# A next generation neural mass model with neuromodulation

**DOI:** 10.1101/2024.06.23.600260

**Authors:** Damien Depannemaecker, Chloé Duprat, Gabriele Casagrande, Marisa Saggio, Anastasios Polykarpos Athanasiadis, Marianna Angiolelli, Carola Sales Carbonell, Huifang Wang, Spase Petkoski, Pierpaolo Sorrentino, Anthony Randal McIntosh, Hiba Sheheitli, Viktor Jirsa

**Affiliations:** Aix Marseille Univ, INSERM, INS, Inst Neurosci Syst, Marseille, France; Department of Engineering, Universitá Campus Bio-Medico di Roma, Rome, Italy; Department of Neurology, University of Minnesota, Minneapolis, MN, United States; Department of Psychiatry and Behavioral Sciences, University of Minnesota, Minneapolis, MN, United States; Simon Fraser University, Vancouver, British Columbia, Canada

## Abstract

The study of brain activity and its function requires the development of computational models alongside experimental investigations to explore different effects of multiple mechanisms at play in the central nervous system. Chemical neuromodulators, such as dopamine, play central roles in regulating the dynamics of neuronal populations. In this work, we propose a modular framework to capture the effects of neuromodulators at the neural mass level. Using this framework, we formulate a specific model for dopamine dynamics affecting D1-type receptors. We detail the dynamical repertoire associated with dopamine concentration evolution and characterize the transitions across qualitatively different oscillatory states as a function of relevant parameters, along with the variations in frequency of the emergent multi-scale oscillations.

## 1 Introduction

Neuromodulators, such as dopamine and serotonin, play central roles in regulating the dynamics of neuronal populations within the central nervous system. These biochemical agents modulate the activities of multiple neurons simultaneously, acting on the global dynamics of the brain.

Neuromodulators can regulate the intrinsic excitability of neurons, influencing their firing rates and patterns of activity [41, 45]. Additionally, at the synaptic level, they can modify the strength of connections between neurons, impacting synaptic transmission and plasticity.

On a broader scale, neuromodulators contribute to the synchronization and coordination of neuronal ensembles, thereby shaping network dynamics involved in various cognitive and behavioral processes [6,18,43,48,52,56]. For instance, dopamine is implicated in reward processing, motivation and motor control [7], while serotonin is associated with mood regulation, sleep-wake cycles, and emotional processing. Dopaminergic pathways have also been implicated in cognitive processes and behaviors such as wakefulness, working memory, and cognitive control [47].

Whole-brain models incorporating the effects of various neuromodulators have been used to investigate the variability in cognitive decline associated with aging [10]. In [31], aging-related changes in dopaminergic regulation have been linked to cognitive performance [5] and hypothesized to play a crucial role in dynamic compensation as a marker of improved cognitive abilities during aging. Impairment in dopamine neuromodulation is also known to play a role in different neurological disorders [29], such as Parkinson’s disease (PD) or Schizophrenia [28].

Psychiatric disorders encompass a heterogeneous group of conditions that disrupt cognition, emotional regulation, and behavior, affecting 970 million people worldwide in 2019, according to the World Health Organization [40]. Among them, schizophrenia impacts approximately 24 million individuals globally. PD and psychiatric disorders are distinct neurological conditions with unique clinical features and prevalence rates. PD, the second most common neurodegenerative disorder, primarily presents with motor symptoms such as tremors, rigidity, and bradykinesia, along with non-motor symptoms. Its prevalence increases with age, affecting 0.04% of individuals aged 40-49 and 2% of those over 80 [4, 42].

In particular, the pathophysiology of PD is characterized by the degeneration of dopamine-producing neurons in the substantia nigra of the basal ganglia, leading to the loss of dopaminergic nigrostriatal neurons along the nigrostriatal pathways, with milder losses in the mesolimbic and mesocortical circuits.

While empirical data provide critical insights into dopamine’s role, computational models are essential for integrating diverse experimental findings, testing mechanistic hypotheses, and predicting emergent neural dynamics that may not be directly observable. Models allow for the systematic manipulation of parameters, simulation of pathological states, and exploration of causal relationships. To model the neuromodulatory aspects of these pathways within personalized whole-brain network models, we need to capture structural changes reflecting synaptic connectivity alterations, particularly within the basal ganglia-thalamocortical circuit [4].

Dopaminergic pathways are also known to play a role in psychiatric disorders via a variety of distinctive mechanisms or dysfunctions in neurotransmission and neuromodulation, primarily featuring the mesolimbic dopamine pathway. This pathway, extending from the ventral tegmental area to limbic regions, takes a central role in the generation of positive psychotic symptoms integral to Schizophrenia. Integrating these diverse neuromodulatory pathways into whole-brain network models will require accounting for inter-regional connection between source and target regions in conjunction with other region-specific parameters. This will be a key ingredient for the development of whole brain network models that can advance our comprehension of the intricate interplay of mechanisms contributing to the emergence of psychiatric and neurological disorders [35, 38].

Furthermore, both PD and psychiatric disorders are accompanied by local and brain-wide changes in neural dynamics. In the context of PD, aberrant dynamics manifest in anomalous bursting activity in the beta frequency range across the basal ganglia-thalamocortical motor network [58], a phenomenon closely associated with clinical disability [57]. Similarly, in psychiatric disorders, there is a disruption in the balance between excitation and inhibition within cortical regions. This imbalance may result from synaptic pruning or alterations in neurotransmitter systems, contributing to the pathophysiology of these conditions [20, 24].

Deep brain stimulation is a therapeutic intervention strategy aimed at rebalancing synchrony dynamics of target neuronal populations in both conditions. Computational modeling studies play a crucial role in providing insight on underlying mechanisms and informing the design of optimal stimulation patterns in silico, thereby advancing the effectiveness of treatment and deepening our understanding of these disorders [17, 33, 51, 59].

Understanding the intricate interplay between neuromodulators action and electrophysiological features of brain is fundamental for elucidating the neural mechanisms underlying complex brain functions and disorders. Mathematical models are indispensable for understanding the workings of multi-scale non-linear interactions out of which emerges the different dynamical repertoires of observed electrophysiological activity.

Additionally, the application of brain network models offers promise of informing the classification and stratification of various forms of these multidimensional neurological conditions. Recent studies underscore this potential of enhancing the classification of PD patients or individuals with psychiatric disorders by supplementing empirical data with simulated data generated from patient-specific brain network models [27].

Integrating personalized whole-brain network models and computational methodologies [19, 55] can contribute to our understanding of the dynamic aspects of these disorders, facilitating refined treatment strategies and a deeper comprehension of their multifaceted etiologies, as it has been done previously for other conditions, such as epilepsy [12]. For that aim, there is a need for a mathematical model that incorporates how neuromodulators influence the electrophysiological activity patterns in the human brain. In order to capture macroscopic brain wide effects and allow for comprehensive parameter exploration, the said model ought to provide a level of description [11,13] able to capture relevant features observed in human data [1,3] in a computationally efficient framework; As such, it will be formulated on the mesoscopic scale as a neural mass model to be implemented in the simulation environment of The Virtual Brain (TVB) [2, 46].

It is crucial to consider the impact of degeneracy and multi-realizability [16] (i.e. multiple models or parametrizations can lead to similar observable behaviors), as well as experimental challenges associated with biophysical mechanisms that are difficult to access, which could impede the interpretability of the model. The selected model should thus incorporate identifiable and measurable parameters that allow for linking biophysical mechanisms to observables of neural dynamics such as known inhibitory and excitatory connections along with accompanying neuromodulation affects on different synaptic receptors activity. It ultimately aims to possess a broad range of dynamic behaviors that is rich enough to encompass complex spatiotemporal features of healthy and aberrant neural activity.

To tackle this challenge, this paper introduces a generic framework for modeling neuromodulator dynamics within a next generation neural mass model [9]. We specifically concentrate on dopamine by presenting a tailored formulation of the corresponding model and an examination of how dopamine dynamics influence neural activity. We present a bifurcation analysis that delineates transitions between qualitatively different oscillatory regimes as the relevant parameters are varied, and provide a quantitative characterization of the variations in the frequencies of emergent multi-scale oscillations in population firing rate.

## 2 Methods

The starting point for this work is a neural-mass model [8], derived by applying a mean-field approach [39] to a network of adaptive quadratic integrate-and-fire model (aQIF) neurons [22]. They key feature of this model, for our purposes, is that it enables the consideration of different conductance-based synaptic inputs.

The aim is to include neuromodulation as a variable in our model. We follow the formalism adopted by Kringelbach et al. [30], which introduces the effects of serotonin on the neural mass level as an additive current, denoted by M, using the Michaelis-Menten equation [25, 36] to capture the dynamics of neuromodulator re-uptake. In our study, we adapt this formalism to account for the generic dynamics of neuromodulators by including the Michaelis-Menten equation for the evolution of neuromodulator concentration, but we depart from [30] by formulating the modulatory effect, *M*, as a multiplicative scaling of the synaptic conductances rather than as an additive current term. This modification aims to enhance the precision and relevance of our model in capturing the nonlinear influence of neuromodulators on neural excitability. Then we propose a specific application for dopamine, with a possible further reduction for this specific case brought about by the relatively slow timescale of dopamine concentration evolution. In the following subsection, we detail the model construction.

### 2.1 Neural mass model

The equations for the neural mass model for the dynamics of a population of *N* neurons are modified version of that studied in the work of Chen and Campbell [8] and Sheheitli and Jirsa [49]. The mean-field derivation invokes the Lorentzian ansatz [39]; assuming a neuronal heterogeneity that takes the form of additive currents, *η*_*i*_, that follow a Lorentzian distribution with a half-width Δ, and centered at 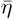, in the thermodynamic limit *N* → ∞, the distribution of the membrane voltages converges to a Lorentzian-shaped function, regardless of the initial conditions. The resulting meanfield variables correspond to the population mean firing rate *r*, mean membrane potential *V* and mean adaptation variable *u*. We consider conductance-based synapses of two types, excitatory AMPA receptors of maximal conductance *g*_*a*_ and reversal potential *E*_*a*_, and inhibitory GABA receptors of maximal conductance *g*_*g*_ and reversal potential *E*_*g*_. This adds two other variables to the model, *S*_*a*_ and *S*_*g*_, for the evolution of the activation of AMPA and GABA synapses. The resulting mean-field equations become:

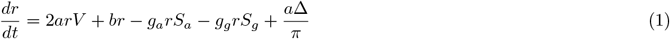

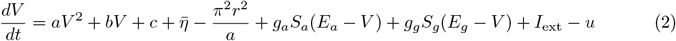

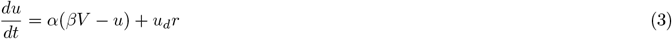

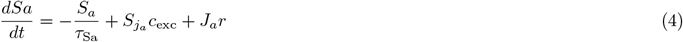

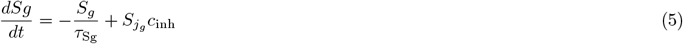

where *a, b, c* are parameters from the original izhikevich formulation of the aQIF [22]; *α, β, u*_*d*_ are respectively, time constant, scaling factor and quantal increment at each spike of the adaptation; 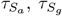 are time constants for AMPA and GABA synapses respectively, *c*_*exc*_, *c*_*inh*_ are external excitatory and inhibitory inputs 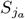 and 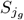 are the post-synaptic strengths of these inputs; *J*_*a*_ is the strength of the recurring connections

This neural mass model set of equations serves as a low-dimensional representation to captures the dynamics of a microscopic neuronal population at the mesoscopic level. We then incorporate in it the dynamics of neuromodulation for the particular case of dopamine.

### 2.2 Generic equations for neuromodulation

At least two aspects are important in neuromodulation dynamics. First, the availability of the neuromodulator is considered, i.e. the evolution of the local concentration. Secondly, its effect on neural activities through specific receptors. One equation for each type of chemical neuromodulation (i.e. capturing the evolution of the concentration) and one equation for each receptor type (i.e. capturing the activation of each of these receptors) are necessary. The functional form for the concentration is made up of two terms, where the first corresponds to the mechanisms leading to an increase of the local concentration which often depend on the activity of a specific brain region that is afferent to the considered node. The second term corresponds to the decrease in concentration due to re-uptake mechanisms that can be captured by Michaelis-Manten formalism. In the following subsection, we describe the construction of the equations specific for dopamine. The variable denoted *M*, captures the proportion of target receptors activated by the neuromodulator. The typical functional form is a sigmoid function [26], the parametrization of which must be specific for each receptor type. If the receptor has an enhancing effect, the slope is positive; and conversely for diminishing effect. Then the *M* variable enters the mean membrane potential equation according to the considered receptor’s specific biophysical properties. It can either modulate a non-synaptic conductance through an additional term of the form *Mg*_*i*_(*E*_*i*_ − *V* ) with *g*_*i*_ and *E*_*i*_ associated with a charged molecule; or directly modulate the synaptic conductances, as it is the case for dopamine. In the following subsection, we take the example of D1 receptors modulating the conductance of AMPA synapses.

#### 2.2.1 Dopamine reuptake

The reuptake of dopamine is a process in which dopamine is returned to the presynaptic neuron from the synaptic cleft. The primary molecular mechanism responsible for dopamine reuptake involves a protein known as the dopamine transporter (DAT). The reuptake process can be described by the following simplified equation:

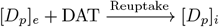

In this equation: [*D*_*p*_]_*e*_ represents dopamine molecules in the extracellular space, DAT represents the dopamine transporter protein on the neuron membrane, and [*D*_*p*_]_*i*_ represents dopamine molecules that have been taken back into the neuron.

The dopamine transporter (DAT) actively transports dopamine from the extracellular space back into the presynaptic neuron, terminating the signal transmission at the synapse. This reuptake process is crucial for regulating the dopamine concentration in the synaptic cleft and maintaining proper neurotransmission. Additionally, it serves as a target for various drugs, including certain antidepressants and psychostimulants, which can modulate dopamine reuptake.

#### 2.2.2 Michaelis-Menten Formalism for Dopamine Reuptake

The dynamics of dopamine reuptake can be described using Michaelis-Menten (M-M) formalism, which involves applying the M-M equation to represent the process of dopamine transport via the dopamine transporter (DAT).

The general form of the M-M equation is:

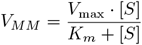

and is a mathematical description of the relationship between substrate concentration [*S*] and reaction rate *V*, characterized by the maximum reaction rate (*V*_*max*_) and the Michaelis constant (*K*_*m*_). We can adapt this equation to describe the dynamics of dopamine reuptake:

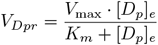

where: *V*_*Dpr*_ is the reuptake rate and represents the velocity dopamine returns to the neuron. *V*_max_ is the maximum reuptake rate, which corresponds to the rate when the dopamine transporter (DAT) is fully saturated with dopamine. *K*_*m*_ is the Michaelis constant, representing the extracellular dopamine concentration at which the reuptake rate is half of *V*_max_. [*D*_*p*_]_*e*_ is the concentration of dopamine in the extracellular space.

This formulation is a simplification, and the actual dynamics of dopamine reuptake are influenced by various other factors, including the number and activity of dopamine transporters, the membrane potential, and the presence of other substances that may modulate reuptake.

It’s important to note that while the M-M formalism provides a useful approximation for enzymatic-like processes, it may not capture all the complexities involved in dopamine dynamics within the synapse. More sophisticated models may be necessary to achieve a more detailed mechanistic understanding of neurotransmitter reuptake kinetics. In our case, we are interested in the global phenomenon leading to the regulation of the electrophysiological activities of a whole population of neurons.

The local concentration increase is driven by input projections from dopaminergic neurons, through the dopaminergic coupling *c*_*dopa*_, scaled in each efferent region by a factor *k*:

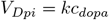

Thus, the change over time (with time constant 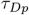) in the extracellular dopamine concentration corresponds to the net result of increase at the rate of local input and decrease at the rate of reuptake:

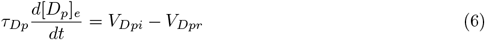

Given the M-M equation for dynamics of dopamine reuptake and the local dopamine concentration, we obtain:

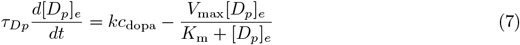

Here, the variation in dopamine concentration occurs on a much slower timescale than that of the evolution of the firing rate dynamics and can be treated as a constant that does not alter the derivation of the mean-field equations associated with the faster variables [8].

#### 2.2.3 Modulatory dynamics for D1 receptors

The following equation represents a mathematical model (inspired from previous work [30]) describing the dynamics of receptors within the populations. It uses a phenomenological activation function and an exponential decrease over time. It can be specifically formulated for the D1-type dopamine receptor.

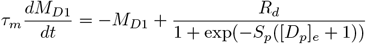

Where: *M*_*D*1_ is the modulation effect associated with the D1-type dopamine receptor. The modulation effect decreases over time (−*M*_*D*1_). *R*_*d*_ is the receptor density in a given receptor population. [*D*_*p*_]_*e*_ is the extracellular dopamine concentration. *S*_*p*_ is a parameter that influences the sensitivity of the receptor to changes in extracellular dopamine concentration. The term 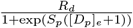 represents the influence of extracellular dopamine on the modulation effect. The sigmoidal function introduces a non-linear activation of the receptor and simulates a saturation effect as dopamine concentration increases.

### 2.3 Neural mass model including neuromodulation

The modeling framework can be adjusted to account for various neuromodulators and receptor types (see Figure 1).We will here focus on dopamine and its corresponding receptors as a representative proof of concept example. Dopamine affects AMPA excitatory conductances [50]. In this case, the *M*_*D*_ variable modulates the conductance *g*_*a*_ through the term *M*_*D*_ + *B*, the variable *M*_*D*_ evolves in a range from 0 to *R*_*d*_ and *B* = 1 is the basal level to enable the minimal AMPA conductance even in the absence of modulation by the extracellular dopamine [*D*_*p*_].

**Figure 1.**
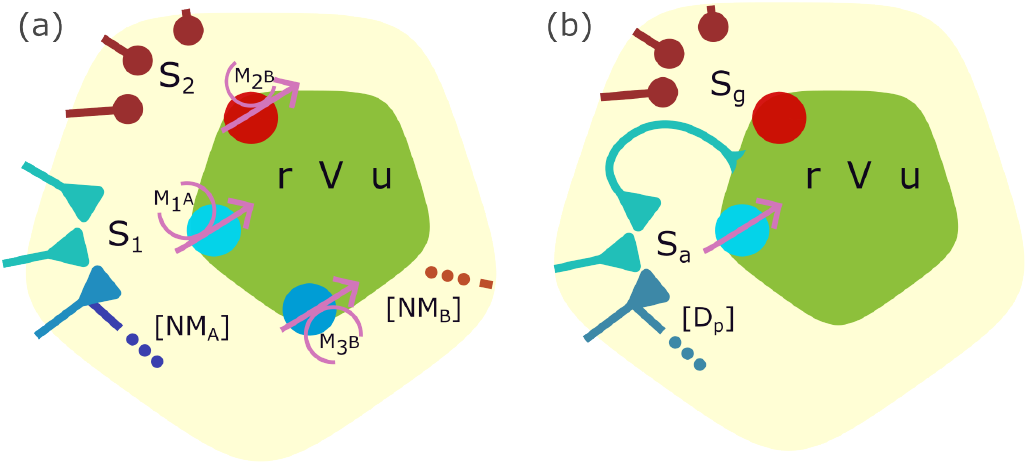
Schematic of the model. (a) A generic neural mass model framework with neuromodulation. Neural activity is characterized by the firing rate *r*, the mean membrane potential *V*, and mean adaptation *u*. It receives different synaptic inputs *S*_1_, *S*_2_, …, and undergoes modulation through receptors *M*_1*A*_, *M*_2*B*_, *M*_3*B*_, … associated with different neuromodulators *NM*_*A*_, *NM*_*B*_, … released by projecting neurons. The modulation can affect either post-synaptic receptors (e.g., *M*_1_ or *M*_2_ in the schematic) or receptors on the neuronal membrane (e.g., *M*_3_). This flexible framework allows for the consideration of multiple neuromodulators and receptors as needed by the scientific question. (b) The model schematic for a neural mass model, as introduced in section 2.3. It can be parameterized for excitatory AMPA and inhibitory GABA synapses, with activation variables *S*_*a*_ and *S*_*g*_, respectively. It also accounts for recurrent excitatory synaptic connections among neurons within the same population. Moreover, it can be tailored for specific cases such as dopamine ([*D*_*p*_]) modulation, which linearly affects AMPA conductance.

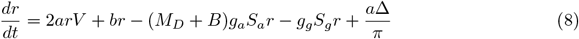

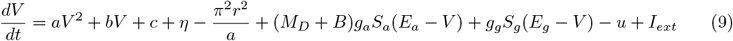

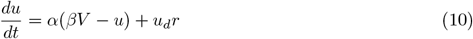

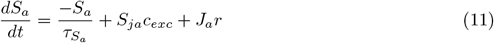

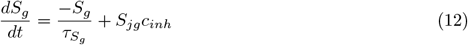

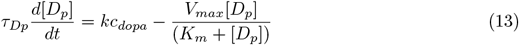

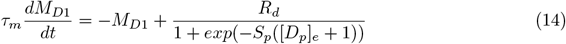

With this formalism, other neuromodulators can be introduced, with an additional variable of the same form as equation 13 for each of them. Then multiple types of receptors can be considered [37], and each of the receptor dynamics would take the same form as the equation 14. Finally, the modulatory equation of each type of receptor would enter the equation 9 of the mean membrane potential according to its biophysical properties and either affect one of the synaptic conductance and/or an additional conductance. The framework is schematized in Figure 1.

In the specific case of dopamine, the time scale of modulation and that of concentration evolution are similar and *M* is strongly correlated with the [*D*_*p*_] variable. In the analysis to follow, we will thus consider only the linear part of the sigmoid of the *M*_*D*1_ equation (specific for D1 receptors), effectively assuming that *M* can be absorbed into the [*D*_*p*_] variable by rescaling. Then, we obtain the linear relation with the form *A*_*D*1_[*D*_*p*_] + *B*_*D*1_, and the equations become:

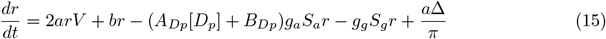

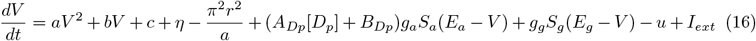

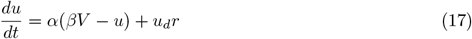

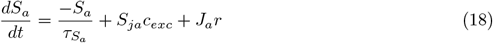

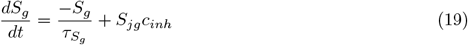

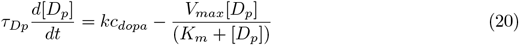

These equations establish our general neural mass modeling framework with neuromodulation. All fixed parameters are given in the Appendix, and are taken from previous work [8, 22] or tuned in a range that insures the dynamical repertoire. We used python for simulation and bifurcation analysis and numerical continuation are performed by exploiting the Julia package BifurcationKit.jl [53].

## 3 Results

We studied the emerging dynamics of this model. We considered an excitatory population, with recurrent connections receiving excitatory and dopaminergic inputs. Given the number of dimensions of the system which obstructs an analytical study of its bifurcation structure, we resort to numerical analysis to unfold the different dynamical regimes that can emerge as the different parameters are varied. Specifically, similarly to previous works [23, 44], we leverage the time scale separation that exists in the system, between the fast sub-system associated with the *r, V* and *S*_*a*_, on one hand, and the slow variable *u* and ultra-slow [*D*_*p*_], on the other. We start with a bifurcation analysis of the fast sub-system constituted by the *r, V, S*_*a*_ variables, considering the slow variables *u* and [*Dp*] as parameters of this subsystem. Then, through numerical simulations, we characterize the full model’s behaviors for different excitatory and dopamine inputs. The parameter values used in the following subsections are given in the Appendix.

### 3.1 Fast-subsystem *r, V, S*_*a*_

In this first subsection, we focused on the sub-system *r, V, S*_*a*_, the three fast variables of the system, revealing diverse qualitative behaviors, notably the first emergence of a limit cycle, and a region of bistability. The variables *r* and *V* were selected as primary observables constituting the fast sub-system. Synaptic activation *S*_*a*_ exhibits nearly the same rapid dynamics, depending on *r*. Importantly, this variable is influenced by dopamine levels in the *r* and *V* equations. Therefore, *r, V* and *S*_*a*_ are the 3 fast variables, while *u* and [*D*_*p*_] being slow and ultra-slow variables, respectively, are considered fixed here (i.e. treated as parameters). Thus, to investigate the dynamics of the fast sub-system we performed bifurcation analysis with respect to parameters *u* and [*D*_*p*_]. *u* has been identified as responsible for bursting behaviors in previous work [8]. Here, bursting refers to population-level activity in spiking neurons, characterized by periods of synchronized firing separated by phases of reduced, asynchronous activity. We chose [*D*_*p*_] as the second bifurcation parameter as we would like to see how the external dopamine concentration affects the behavior of our neural population. The result of the analysis is shown in 2. In the chosen range of parameters, we observe the emergence of a region, in white in Fig. 2 I, with a single stable fixed-point (Fig. 2 II (A)). In the light blue region, delimited by a Hopf curve, the fixed point loses stability in favor of an attracting limit-cycle, leading to the oscillations shown in Fig. 2 II (B). For higher values of [*D*_*p*_] and/or *u*, after crossing again the Hopf curve, the system returns to a fixed point, associated with saturation, at a low firing rate *r* and high value of mean membrane potential *V*, as shown in Fig. 2 II (C). Two curves of fold (saddle-node) bifurcations delimit a region, shaded in orange, within which the fast sub-system presents a bistable behavior, characterized by the co-existence of an up and a down state. An example of the dynamics of the system for parameters value inside this region is represented in 2d, which shows the co-existence of a state of higher activity, characterized also by higher mean membrane potential, with another state of low firing rate, with the value of V close to the resting membrane potential. Finally, the overlap between the orange and blue regions is characterized by the bistability between a fixed point and a limit cycle, as observed in Fig. 2 II (E).

**Figure 2.**
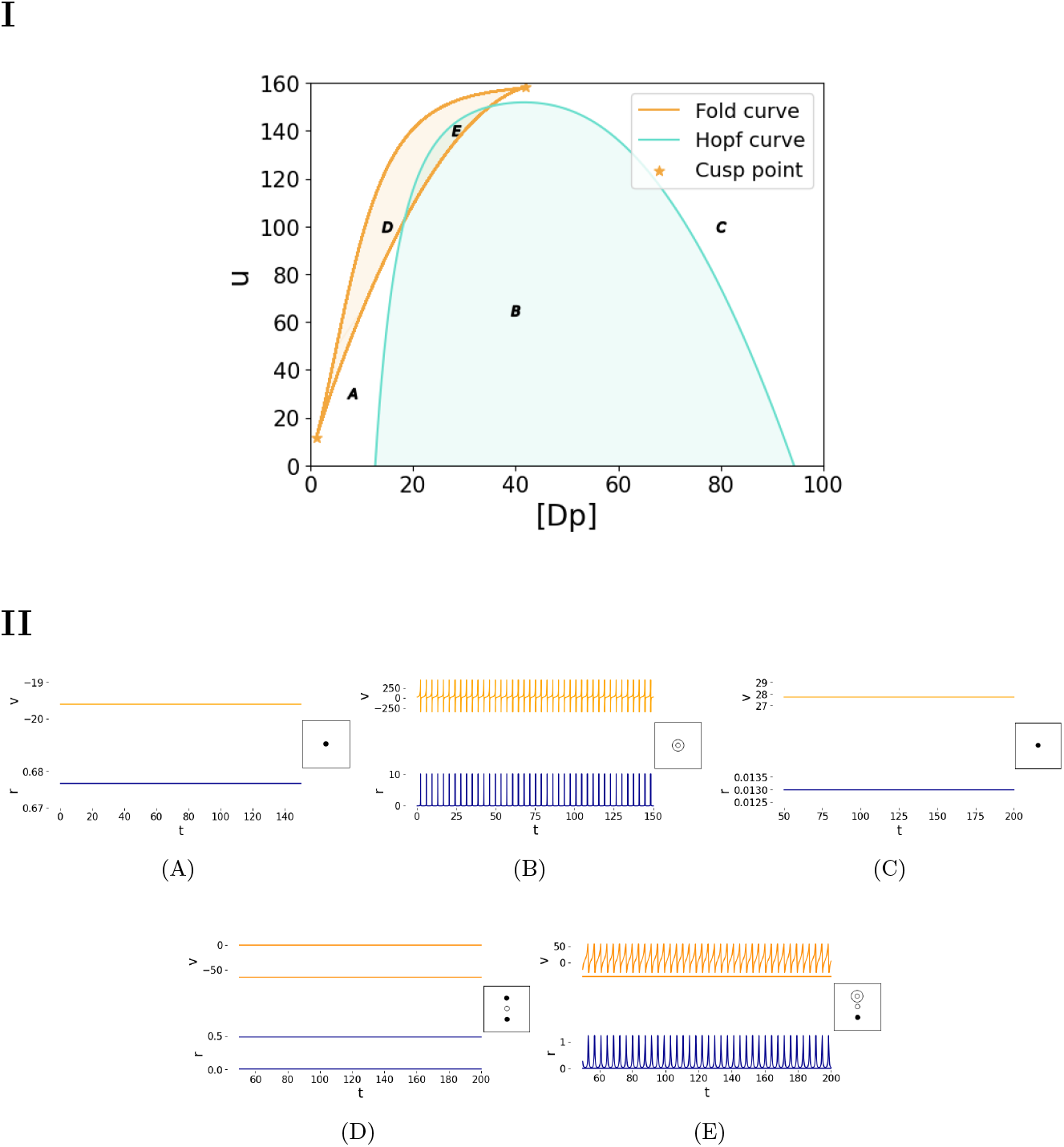
Dynamical patterns of the fast sub-system: I. Bifurcation diagram of the fast sub-system (*r, V*, *S*_*a*_) with respect to *D*_*p*_ and *u*, and the corresponding time series for each *Dp*. Each time series is associated with a sketch of the phase portrait, where black (white) dots represent stable (unstable) fixed point and solid black circle represents stable limit-cycles. II. (A) fixed-point behavior of high activity (*r >* 0.5, (B) oscillation associated with a limit-cycle corresponding to synchronious activity, (C) fixed-point behavior of low activity (*r <* 0.05, (D) bistable fixed-point behaviors, (E) bistability between limit-cycle and fixed-point.

### 3.2 Full system dynamics

After having described the possible dynamics of the fast *r, V, S*_*a*_ subsystem, we present the dynamical repertoire of the full system of equations. The two slower variables *u* and [*D*_*p*_], enable transitions between the qualitatively different behaviors described in the previous subsection.

In this section, we present the results of numerical simulations of the model for different values of external inputs, *c*_*dopa*_ and *c*_*exc*_, with different values of the parameter *u*_*d*_ associated with adaptation mechanisms and different cell types [22]. The rationale behind choosing these three parameters is that this neural mass model is built to be used in whole-brain models [3], where the nodes representing different brain regions (possibly associated with different cell properties capture by *u*_*d*_), are interconnected through dopaminergic *c*_*dopa*_ and long-range excitatory *c*_*exc*_ projections. We performed simulations using numerical integration, allowing the complete system to evolve for 10 seconds and, then eliminating the transient period of 6.5 seconds, after which the system stabilizes into a steady-state; we not only focus on the mean firing rate of the steady state but also on the standard deviation to identify parameter values for which the dynamics undergo significant changes in amplitude variability that is indicative of oscillatory behavior. These metrics are presented in Figure 3, where various qualitative behaviors are illustrated, as also depicted in Figure 4. Specifically, fixing the excitation and adaptation parameters, ( *c*_*exc*_ = 0.2, *u*_*d*_ = 12), it is evident that as *c*_*dopa*_ increases, the system undergoes transitions in its dynamical behaviour, reflected in changes in both the average firing rate and its standard deviation.

**Figure 3.**
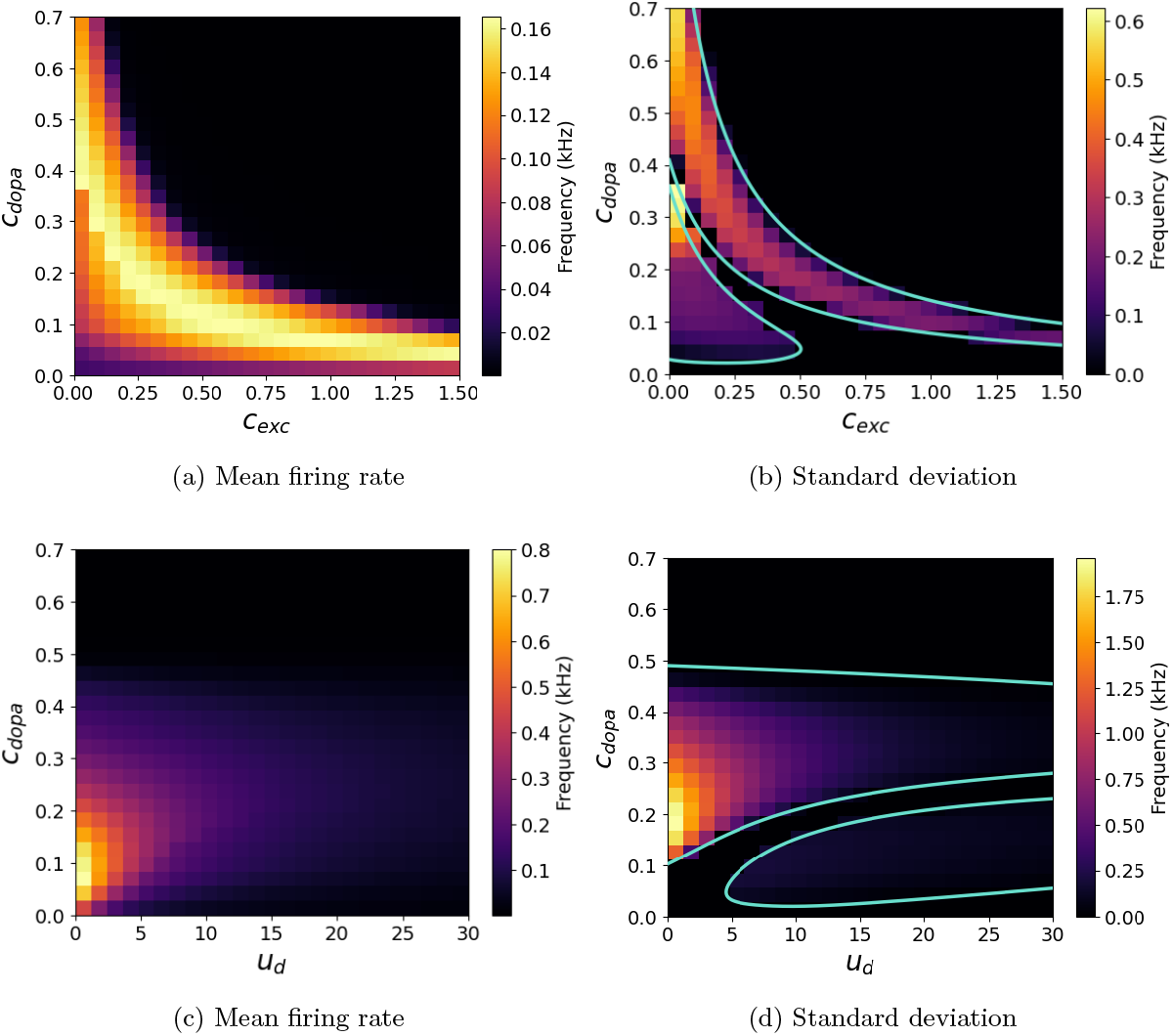
After transient behavior of the complete system for different values of parameters. In the first row, effects of excitatory and dopaminergic inputs on a single neural node, with the mean firing rate shown on the left (a) and its standard deviation on the right (b). The value of the parameter *u*_*dopa*_ = 12 is fixed. In the pictures, the colour gradient transitions from purple to yellow, marking the shift from the lowest to the highest values observed. The highest mean firing rates occur mostly for low value of *c*_*dopa*_. Non-zero standard deviation indicate oscillatory behaviors. The light-blue lines indicate the bifurcation line found using continuation tools. Below the same analysis is shown for different effects of dopaminergic and adaptation inputs, in this case *c*_*exc*_ = 0.2 is constant. In general mean firing rate (c) and its standard deviation (d) have larger values if compared to the previous case. However, once again, high values for the mean firing rate are observed for low dopaminergetic inputs. Non-zero standard deviation indicate oscillatory behaviors.

**Figure 4.**
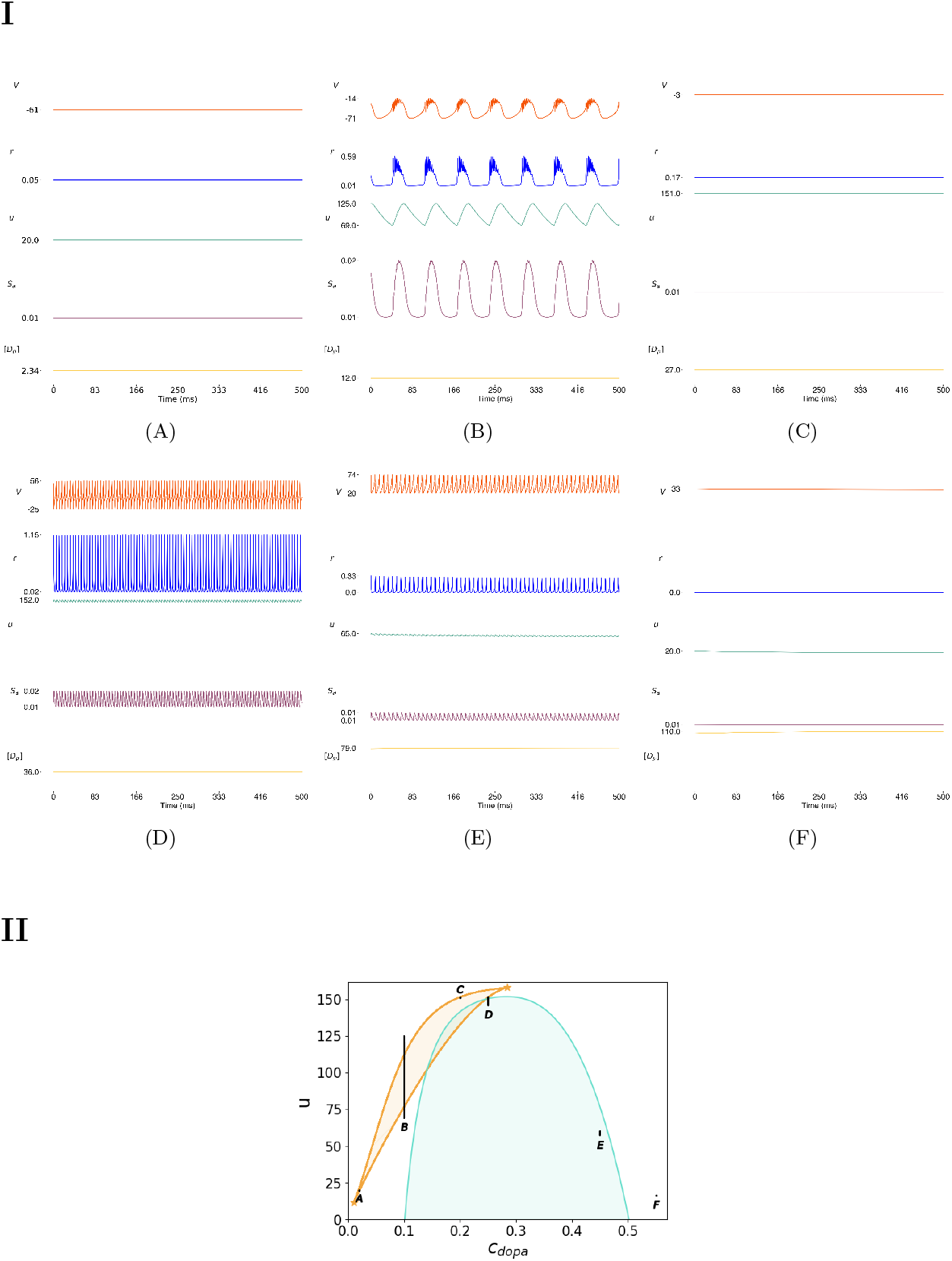
Dynamics of the full system : (I) Time series of distinct variables of the full system for increasing values of *c*_*dopa*_, at fixed *c*_*exc*_ = 0.2 and *u*_*d*_ = 12. (A) Low values of *c*_*dopa*_ (∼ 0.02) are characterized by a stable fixed point. (B) Increasing the dopaminergic input (∼ 0.1) one notices the emergence of bursting behavior. (C) A further increment in *c*_*dopa*_ ( ∼0.2) makes the system transits to stable behaviour before acceding again an oscillatory region (D) with higher frequency with respect to the previous one (*c*_*dopa*_ ∼0.25). (E) Further on (∼ 0.45) the system remains in oscillatory regime but gradually decreasing both oscillations amplitude an frequency. (F) For higher values of *c*_*dopa*_ ( ≥ 0.55) the full system presents a constant stable fixed point, although with different values with respect to the initial low dopaminergic one. All simulations were conducted with “resting state” conditions with *r*_*init*_ = 0.01 kHz, *V*_*init*_ = -70 mV, *u*_*init*_ = 0, 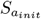 = 0.006, *S*_*g*,*init*_ = 0, [*D*_*p*_]_*init*_ = 0.6. (II) Superposition of time series of the slow variable u and [*D*_*p*_] to the codimension-2 bifurcation diagram of the fast subsystem. In the abscissa we substitute the steady state [*D*_*p*_] with the corresponding value of *c*_*dopa*_, since there exist a one-to-one relation when [*D*_*p*_] reached its steady state.

The dopamine input *c*_*dopa*_ influences the excitability if the neuronal population. In Fig. 7, we numerically evaluate the value of *c*_*dopa*_ at which the system departs from the resting state characterized by low steaty state values of *V* (∼ −64) and *r* (*<* 0.004).

We here analyze the transient behavior of the neural mass, focusing on excitatory, dopaminergic, and adaptive inputs. Our findings highlight how the mean firing rate and standard deviation of the system change under these parameters vary (see Fig. 3). We observe that high mean firing rates occur when excitatory or dopaminergic inputs are low. Non-zero standard deviations indicate the presence of oscillatory behavior. We use continuation tools [53] to identity the Hopf bifurcation lines (light-blue) in Fig. 3 (b) and (d), which mark the boundary where the system changes behavior from fixed-point to limit cycle or bursting behavior. When analyzing dopaminergic and adaptive inputs together, we find generally higher values for both metrics, with high mean firing rates at low dopaminergic input levels while maintaining oscillatory patterns.

This is illustrated in Figure 4 where we present time series of several variables of the full system, for increasing values of *c*_*dopa*_, together with the path defined by the slow variables in the codimension-2 bifurcation diagram, when considering the evolution of the full system. In Figure 4.I, one notices that, starting from a low value of dopaminergetic input, the system shifts from a fixed point to oscillatory behavior (Fig. 4.I-A and Fig. 4.I-B), coinciding with an increment of the value of standard deviation. With further increase in the value of *c*_*dopa*_, the system approaches the separation line for the values of standard deviation visible in Figure 3. Hence, it transitions briefly to stable fixed behavior, before becoming oscillatory again, with a higher frequency (Fig. 4.I-C and Fig. 4.I-D). With an additional increment in the dopaminergic parameter, both the amplitude and frequency of oscillations decrease (Fig. 4.I-E) until reaching another separation line, as depicted in Figure 3. Beyond this boundary, the dynamics settle into a stable fixed point, similar to the scenario at lower *c*_*dopa*_, albeit at different values for each variable of interest (Fig. 4.I-F). In Figure 4.II, we observe how transitions among the different dynamical behaviors for the full system are possible as the “slow” variables are allowed to evolve in time. Here, black lines and dots represent time series of u and [*D*_*p*_] superimposed to the codimension-2 bifurcation diagram of the fast sub-system that was previously presented. We point out that, compared to [*D*_*p*_], which can just exhibit relaxation towards a stable equilibrium, *u* can also display oscillatory behavior due to feedback from the membrane potential 14 variable (a hysteresis effect [44]). This enables the appearance of bursting activity, as presented in Figure4.I-B, where the oscillation of *u* drives the fast sub-system to switch from down to up state and vice versa.

Moreover, we emphasize the fact that, for each value of *c*_*dopa*_, keeping the other parameters in 7 constant, the dopamine concentration reaches a different steady-state value. Thus, in 4.II, we substitute the values of [*D*_*p*_] with the corresponding *c*_*dopa*_ value.

This demonstrates the dopaminergic effect on the neural mass model, how changes in the steady state dopamine concentration value can induce qualitatively different dynamical regimes in the system. We now proceed to further characterize the oscillatory regimes highlighted in Figure 4.

More specifically, we quantify the variations in the frequency of oscillation in those specific regimes. In order to do so, we simulate 8000 ms long time series for different pairs of parameter values inside each region. Then, we considered just the last 3000 ms to avoid transient effects and, depending on the identified dynamical regime, we compute the single or multiple periods of oscillation and obtain the frequency by inverting it.

For the regions displaying fast oscillating behaviour, we simply compute the period as the mean time distance between two consecutive peaks. Results for those are represented in Figure 5a in the (*c*_*exc*_, *c*_*dopa*_) plane, and in Figure 5b for the (*u*_*d*_, *c*_*dopa*_). Both plots show overall decreasing frequency values when the parameter other than the dopaminergic one is increased. Moreover, we notice that the range of frequencies obtained in the (*u*_*d*_, *c*_*dopa*_) plane is much higher than those in the (*c*_*exc*_, *c*_*dopa*_) plane.

**Figure 5.**
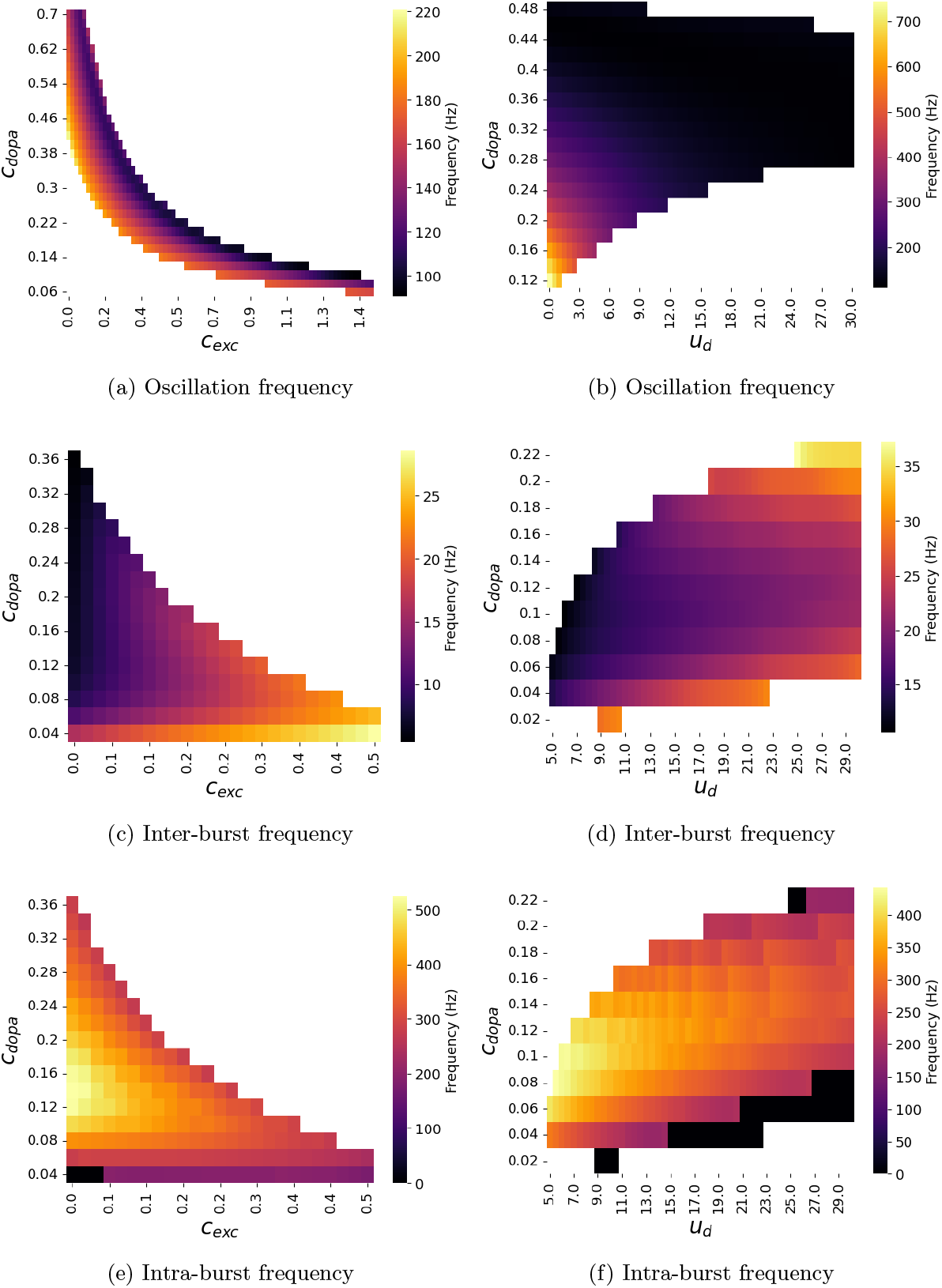
Fequencies observed for the different oscillatory regimes: (a) Oscillation frequencies for different values of *c*_*exc*_ and *c*_*dopa*_ (b) Oscillation fequencies for different values of *u*_*d*_ and *c*_*dopa*_, (c) Slow frequency components of the bursting behavior, i.e. inter-burst for *c*_*exc*_ and *c*_*dopa*_, (d) similarly for *u*_*d*_ and *c*_*dopa*_, (e) and (f) are the corresponding fast frequency component of the intraburst

In the regions characterized by bursting behavior, we identified the peaks of each single burst event by imposing a threshold on the time difference between two consecutive peaks (18 ms) to be assigned as belonging to the same burst event. Then, we compute inter-burst period as the mean time distance between the first peak of each burst, and the intra-burst period by averaging over all the time distances of peaks within the same burst. For this analysis the results are presented in Figures 5c and 5e, for (*c*_*exc*_, *c*_*dopa*_), and in Figures 5d and 5f for (*u*_*d*_, *c*_*dopa*_). We observe an inverse trend for inter-burst and intra-burst frequencies between the different realizations, in the sense that as the former decreases, the latter instead increases. The observed non-linear relationship highlights the complexity of dopaminergic neuromodulation, as it influences electrophysiological activity based on both the excitatory input *c*_*exc*_ and intrinsic cellular properties represented by *u*_*d*_.

Since this neural mass is designed to be used in whole-brain modeling, the dopaminergic projection through *c*_*dopa*_, would become dynamical and enable the transition between the different patterns. We gave examples of these transition by showing in Fig. 6, the transition from fixed-point to bursting behavior (Fig. 6 (a)), from bursting to fixed point (Fig. 6 (b)), from fixed-point to oscillations (Fig. 6 (c)), from oscillations to fixed-point (Fig. 6 (d)) and from bursting behavior to oscillation (Fig. 6 (e)). This is obtained thanks to the transient from the initial condition to steady-state of the the variable [*D*_*p*_] for different constant values of *c*_*dopa*_, parameter values are given in the caption of the Fig. 6.

**Figure 6.**
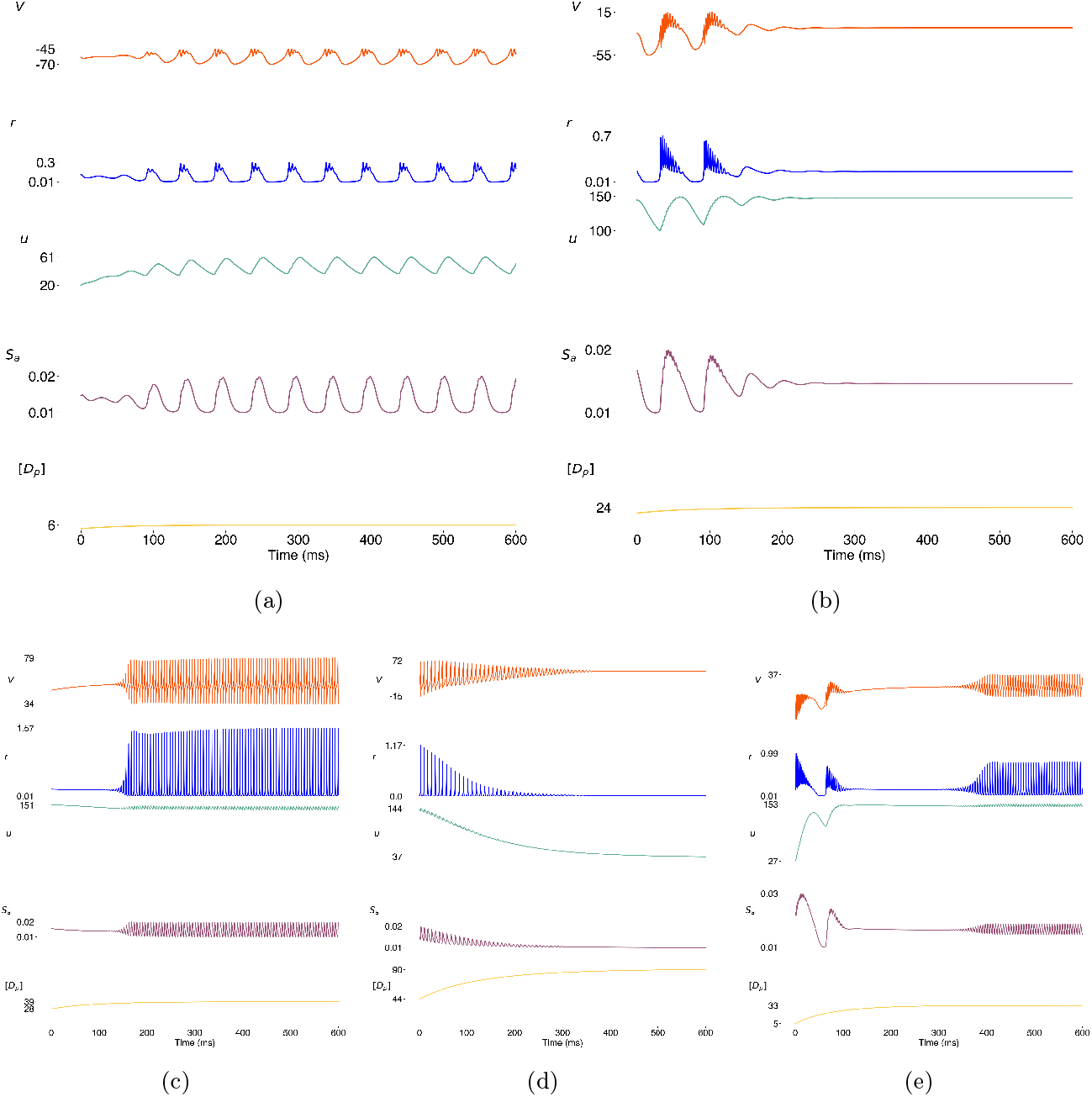
Qualitative transition time series of distinct variables of the full system for increasing values of *c*_*dopa*_, at fixed *c*_*exc*_ = 0.2 and *u*_*d*_ = 12. The ultra slow variable [*Dp*] evolves slowly from the initial condition to steady-state inducing pattern transitions in the rest of the system. In the Sub-figure (6a) shows the system entering a bursting behavior for *c*_*dopa*_ (∼ 0.05), which it then leaves for *c*_*dopa*_ ( ∼0.18) to enter a constant behavior defined by a fixed point (6b). Increasing the dopaminergic input to enter the range of oscillatory trajectories, the system enters ( ∼0.27, 6c) then exits ( ∼0.49, 6d) an oscillatory behavior for a stable behaviour. Sub-figure (6e) shows an interesting behaviour as the full system goes from bursting to stable behaviour and finally to oscillations. All simulations presented in this figure were conducted using “resting state” conditions as initial conditions with *r*_*init*_ = 0.01 kHz, *V*_*init*_ = -70 mV, u_init_ = 0, 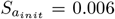, [*D*_*p*_]_*init*_ = 0.6 and transient periods from 0ms to 200ms were removed to focus on the period of interest.

## 4 Discussion

We proposed a modeling framework to capture the effects of neuromodulator dynamics on the neuronal electrophysiological activity at a neural mass level. We presented a reduced version, specific for dopamine dynamics, as a proof of concept, and provided a thorough bifurcation analysis of the qualitatively different possible dynamical regimes that emerge as steady state dopamine concentration and input excitation levels are varied.

Such an approach enables us to capture a rich and complex dynamical repertoire with a reduced number of variables that can still be interpreted or associated with meaningful biophysical units. Indeed, while the proposed model is phenomenological in nature by its high level of description, it can nevertheless serve to explicitly represent and capture the evolution of physical variables that can be directly related to observables, such as firing rates, mean membrane potentials, and synaptic currents. Thus, this is a biophysically inspired model [19], in which the fast dynamics is obtained through mean field theory and complemented with slow dynamics in a heuristic efficient formulation at the mesoscopic scale. The end result is a minimal mechanistic neural mass model that lends itself to parameter fitting for capturing complex electrophisiological oscillatory dynamics that may arise in different types of neuronal populations across brain regions.

While fitting the proposed model to specific experimental data is beyond the scope of the current work, it is worth noting that the frequencies of inter and intra-burst oscillations, emerging for the parameter values studied here, are in the range of what has been observed in basal ganglia neuronal activity recorded from implanted deep brain stimulation electrodes in PD patients [34]. More specifically, the administration of dopamine replacement medication (L-DOPA) was reported to induce a shift in the peaks of the power spectrum of the subthalamic nucleus activity, from 15HZ and 250Hz to 25Hz and 350Hz (see Fig. 2 in [34]). This is inline with what is suggested by the results in Fig. 5e and 5c, that is, there exists a range of parameter combinations for which the proposed model would exhibit inter and intra-burst frequencies that would be consistent with those experimentally reported for the concurrent shift in spectral peaks. We see this as as an encouraging testament to the relevance of the proposed model, particularly in how it pertains to dopamine action in the basal ganglia circuit and the emergence of multi-frequency complex oscillations therein.

One novelty of the framework presented here, in comparison to other neural mass modeling approaches, is the expression of the action of dopamine as a scaling of the synaptic conductance at the mesoscopic neural mass model level; this formulation can in fact be seen as a natural extension of the seminal works of Humphries [21] and Durstewitz [14, 15] in which dopaminergic action was modeled as such but on the microscopic level of individual spiking neurons. This latter approach was also employed in [32] to capture the effects of dopamine depletion in a multi-population spiking neural network model of the basal ganglia. As such, our proposed form for dopamine action on the neural mass model builds on previous work of validated biophysical models of spiking neurons to provide a computationally efficient framework that extends the modeling of dopaminergic neuromodulatory action, from the level of populations of spiking neurons, to that of the whole brain network level.

This modeling framework also enables the introduction of effects on other receptors, such as NMDA type receptors [49]. The framework is modular and can be adapted to fit the scope of different applications; that is, it can serve as a foundational building block to be used in personalized brain models, such as in the context of virtual brain twins [55]. Virtual brain twins is a novel concept, in which only features that are of interest to a neurological condition are selected to be captured by the model, so as to maximize efficiency and relevance to corresponding clinical data [55]. This methodology aims to then personalize the model to the individual patient’s brain by constraining the model to anatomical and funcitonal patient-specific data using advanced parameter fitting algorithms. The resulting models can then serve as in-silico simulation engines for the guidance of more optimal diagnosis and treatment of brain disorders. By tailoring the modeling to individual subjects, researchers can better understand the intricacies of brain function and mechanisms of its pathology. In Wang et al. [55], the standard model was presented in a generic format, including local dynamics with both global and local connectivity. With this work, we present a first detailed neural mass model that allows the incorporation of neuromodulator dynamics on a mesoscopic level in large-scale whole-brain dynamics. Ultimately, the proposed model paves the way for the development of virtual brain twins [54], particularly in the context of PD [3] and psychiatric disorders, for which the disruption in chemical neuromodulator action is known to play a significant role in the underlying psychophysiology.

## 5 Acknowledgment

The preparation of this article was funded through the EU’s Horizon Europe Programme SGA 101147319 (EBRAINS 2.0), and SGA 101137289 (Virtual Brain Twin). The project leading to this publication has received funding from the Excellence Initiative of Aix-Marseille Université - AMidex, a French “Investissements d’Avenir programme” AMX-21-IET-017. We acknowledge the support of the Government of Canada’s New Frontiers in Research Fund (NFRF). The project leading to this publication has received funding from the Excellence Initiative of Aix-MarseilleUniversité - A*Midex, a French “Investissements d’Avenir programme” AMX-21-IET-017.

## APPENDIX

### parameters values

In the following table, we detail the parameter values used to generate the presented results.

|  | Variables | Symbol | Associated unit |
| --- | --- | --- | --- |
| Neuromodulation<br>neural mass | Firing rate | $r$ | kHz |
| | Mean membrane potential | $V$ | mV |
| | Adaptation | $u$ | nA |
| | AMPA synapses activation | $S_a$ | - |
| | GABA synapses activation | $S_g$ | - |
| | Extracellular dopamine concentration | $[D_p]_e$ | mM |
| | D1 dopamine receptor modulation | $M_{D1}$ | - |

| Parameters | Symbol | Value | Associated unit |
| --- | --- | --- | --- |
| aQIF parameter | $a$ | 0,04 | - |
| aQIF parameter | $b$ | 5 | - |
| aQIF parameter | $c$ | 140 | - |
| aQIF parameter | $\alpha$ | 0,013 | - |
| aQIF parameter | $\beta$ | 0,4 | - |
| Mean additive current | $\eta$ | 18 | nA |
| Maximal conductance of AMPA | $g_a$ | 12 | nS |
| Maximal conductance of GABA | $g_g$ | 12 | nS |
| Additive current half-width distribution | $\Delta$ | 1 | - |
| Linearization factor of dopaminergic receptors | $A_{D_p}$ | 1 | nS/mM |
| Basal level in absence of neuromodulation | $B$ | 0,2 | nS |
| Reversal potential of AMPA synapses | $E_a$ | 0 | mV |
| Reversal potential of GABA synapses | $E_g$ | -80 | mV |
| External current | $I_{ext}$ | 0 | nA |
| Adaptation step | $u_d$ | 12 | nA |
| Time constant of AMPA synapses | $\tau_{S_a}$ | 5 | ms |
| Time constant of GABA synapses | $\tau_{S_g}$ | 5 | ms |
| Time constant of dopamine concentration evolution | $\tau_{D_p}$ | 500 | ms |
| Time constant of dopamine receptor modulation | $\tau_m$ | 500 | ms |
| Strength of AMPA synapses | $S_{j;a}$ | 0,8 | nS |
| Strength of GABA synapses | $S_{j,g}$ | 1,2 | nS |
| Excitatory coupling | $c_{exc}$ | - | kHz |
| Inhitatory coupling | $c_{inh}$ | - | kHz |
| Dopamine coupling | $c_{dopa}$ | - | mM |
| Maximum dopamine reuptake rate | $V_{max}$ | 1300 | mM/s |
| Michaelis constant | $K_m$ | 150 | mM |
| Dopamine coupling factor | $k$ | 100 000 | - |
| Receptor density | $R_d$ | 1 | - |
| Receptor sensitivity to dopamine | $S_p$ | 1 | - |
| Noise standard deviation | $\sigma$ | $1e - 3$ | - |

**Figure 7.**
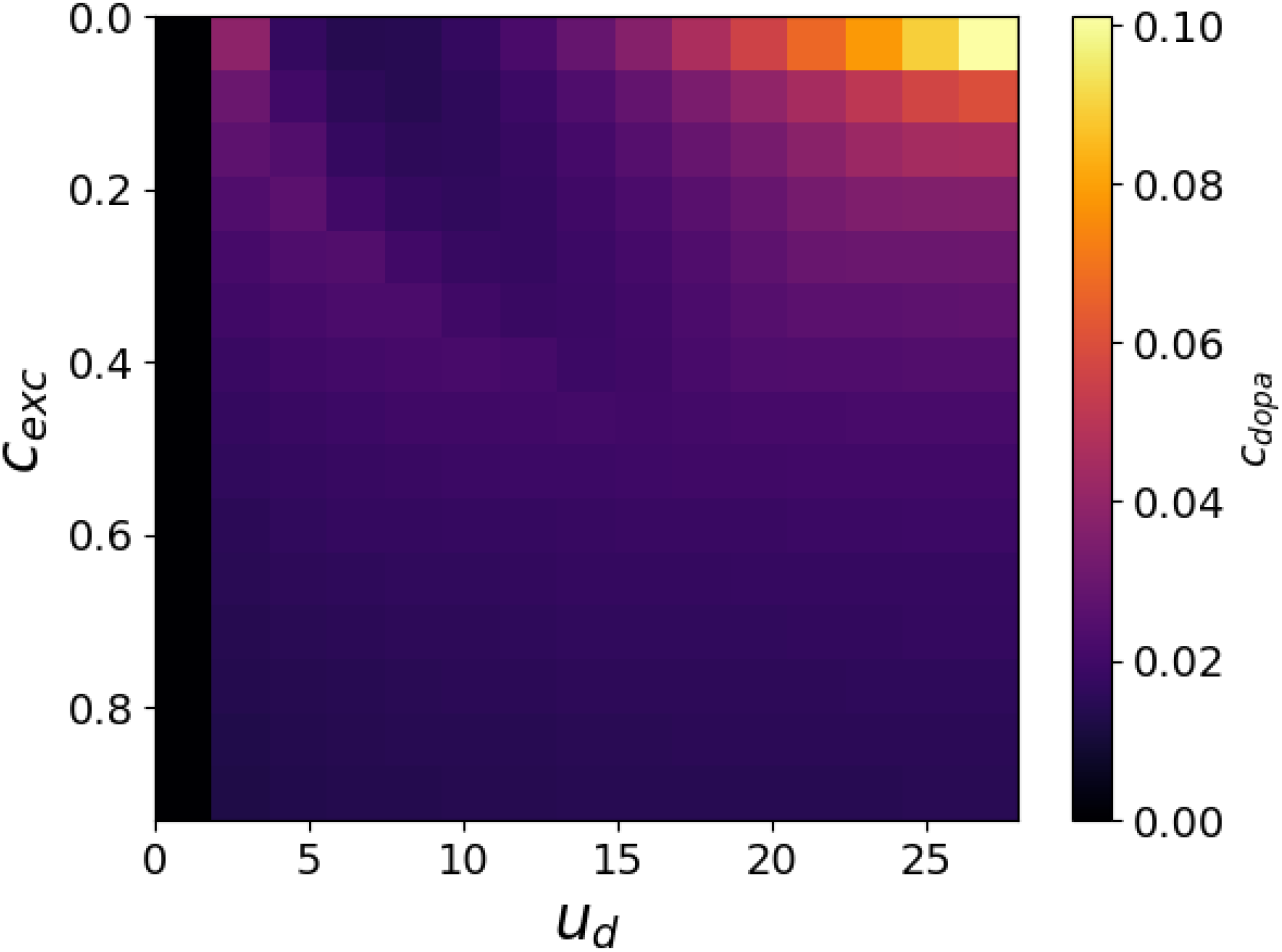
*c*_*dopa*_ value for which the system exit the resting state (RS), for different values of the parameters *u*_*d*_ and constant external input *c*_*exc*_

